# Chronic Subconvulsive Activity during Early Postnatal Life Produces Autistic Behavior in the Absence of Neurotoxicity in the Juvenile Weanling Period

**DOI:** 10.1101/645705

**Authors:** LK Friedman, BA Kahen

## Abstract

The diagnosis of autism spectrum disorder (ASD) varies from very mild to severe social and cognitive impairments. We hypothesized that epigenetic subconvulsive activity in early postnatal life may contribute to the development of autistic behavior in a sex-related manner. Low doses of kainic acid (KA) (25-100 µg) were administered to rat pups for 15 days beginning on postnatal (P) day 6 to chronically elevate neuronal activity. A battery of classical and novel behavioral tests was used, and sex differences were observed. Our novel open handling test revealed that ASD males nose poked more often and ASD females climbed and escaped more frequently with age. In the social interaction test, ASD males were less social than ASD females who were more anxious in handling and elevated plus maze (EPM) tasks. To evaluate group dynamics, sibling and non-sibling control and experimental animals explored 3 different shaped novel social environments. Control pups huddled quickly and more frequently in all environments whether they socialized with littermates or non-siblings compared to ASD groups. Non-sibling ASD pups were erratic and huddled in smaller groups. In the object recognition test, only ASD males spent less time with the novel object compared to control pups. Data suggest that chronic subconvulsive activity in early postnatal life leads to an ASD phenotype in the absence of cell death. Males were more susceptible to developing asocial behaviors and cognitive pathologies, whereas females were prone to higher levels of hyperactivity and anxiety, validating our postnatal ASD model apparent in the pre-juvenile period.

**Highlights:**

- Chronic subconvulsive activity in early life leads to autism phenotypes.
- Juvenile males were susceptible to asocial behaviors and cognitive pathologies.
- Juvenile females were prone to hyperactivity and anxiety validating sex differences.
- Non-siblings were erratic in groups irrespective of sex.
- A postnatal epigenetic model may drug screen for milder forms of autism.

## 1. INTRODUCTION

Autism spectrum disorder (ASD) is an early childhood neurodevelopmental disorder of varying severity ranging from mild to severe which includes asocial behavior, cognitive deficits, and repetitive stereotypic behaviors [1]. Despite the high prevalence of ASD in young children, the etiology of autism remains unknown, as is the higher incidence in males [2, 3]. From 2000-2010, epidemiology studies showed that the prevalence of ASD in the US increased by 331% with a higher incidence in males carrying a genetic mutation (e.g. the 22q11.2 duplication), however, a higher prevalence was also found in females carrying rare copy-number variants [4–6]. More recent reporting reflects a further increase from 2004 to 2014 in ASD prevalence and incidence was observed based on sex, region, and socio-economic level, possibly due to heightened awareness of variations of ASD [7]. Milder forms of autism have a later developmental onset compared more severe cases (18 months vs. 34-48 months) [8]. Genetic disorders such as fragile-X mental retardation protein (FMRP) gene deletion are a well-studied model of ASD in rodents since a fragile X mutation is the most common genetic cause of symptoms. However, it is important to note that this gene only accounts for 2-3% of all ASD cases so that other populations without this genetic defect are not represented [9]. Similarly, mutations of other genes such as SCN1, SCN2A, GABRA1, GRIN1, GRIN2B result in more severe ASD phenotypes whereby seizures are a co-morbidity [10, 11], but again these genes do not account for the mild and more functional ASD cases that are not pervasive until years after birth and do not exhibit seizures [12]. Since prenatal exposure to valproic acid (VPA) is currently used to treat epilepsy and Bipolar Disorder [13, 14], which can lead to an ASD phenotype [15, 16] administration of VPA to pregnant female rats has been extensively used as an animal model of autism with prenatal factors [13, 14, 17, 18]. However, once again the percentage of children diagnosed with ASD prenatally exposed to VPA is relatively low (4.1% for ASD + epilepsy and 2.9% for ASD) [19, 20]. Therefore, preclinical immature rodent models are equally needed to address environmental postnatal stress factors that can lead to autistic pathology.

Clinical outcomes of later onset disorders are likely driven by early postnatal events that can shape maturation of the central nervous system (CNS), such as apoptosis, neurogenesis, synaptogenesis, synapse elimination, dendritic arborization and pruning as well as the excitatory/inhibitory balance [21]. For example, changes in the excitatory/inhibitory balance by chronic subseizure levels of glutamate may underlie core ASD symptoms via long-lasting or permanent defects in non-ionotropic receptors such as phosphorylation of cannabinoid receptors (CBR) and elevations in endocannabinoid (eCB) content [22, 23]. Accordingly, in the prenatal valproic acid (VPA) autism model, deficits in anandamide (AEA) metabolism and phosphorylation of CB1 receptors were observed in the hippocampus and amygdala in the developing and matured rat which may contribute to behavioral pathology [24]. Administration of VPA to pregnant female rats has been extensively used as a model of ASD with prenatal factors [17, 18, 25] since clinically it was found to be a teratogen and danger to pregnant women by increasing the incidence of ASD in the offspring [13, 14]. In contrast, we theorized that postnatal stress factors may also elicit mild autistic traits that are different from children with epilepsy having an ASD phenotype [26]. Early life seizure models have revealed ASD traits in adulthood following long-lasting morphological and physiological changes as a consequence of recurrent early life convulsive seizures elicited by 50-75 flurothyl [27, 28]. Milder forms of ASD not associated with seizures may be distinguishable and more treatable if detected at pre-juvenile ages.

Because early detection is critical to obtain clinical intervention that improves cognitive and adaptive behavior in toddlers diagnosed with ASD [29, 30], this study focused on a preclinical immature subconvulsive activity rat model using micromolar doses of a glutamate analogue, kainic acid (KA) to produce reproducible behavioral change. Hence, our prior research using KA-induced status epilepticus convulsive model was modified to a chronic subconvulsive postnatal model assumed to cause elevations in basal glutamate but without triggering seizures. This perinatal KA dosing model is consistent with a previously established rodent model of ASD that used low (subconvulsive) doses of demoic acid (DOM) or KA, but their paradigm was initiated in the second postnatal week, which resulted in long-term changes in behavior and histopathology that manifested in adulthood [31]. In a more recent study, DOM was exposed prenatally to mice which caused sex-related behavioral changes that resembled diagnostic features relevant to ASD in males [32, 33]. Alterations in sociability, social recognition, and stereotypic behaviors generally have been assessed in adulthood [34]. We questioned whether chronic but subtle elevations in postnatal glutamate levels can cause similar effects to induce autistic / intellectual disability and how long it takes to cause mild forms of autistic pathology to manifest in the immature brain. Another goal was to examine the initial onset of sex-related behavioral change which may facilitate future drug screening during a critical period of development that is associated with regulated surges of stress hormones. Therefore, subconvulsive treatment began in the first postnatal week and animal behavior was tested in the pre-juvenile weanling period to optimize an early window of therapeutic intervention [35]. Both classical and non-classical behavioral testing were utilized.

## 2. METHODS

### 2.1 Animals and Drug Administration

To develop a reliable subconvulsive model for autism in development, male and female mixed and same sex litter rat pups were randomly assigned to receive three subconvulsive doses of KA for 15 days beginning on P6 with three 5-day step-up intervals. Kainic acid (Cayman Chemicals) was dissolved in phosphate buffered saline (PBS) to a stock solution of 10 mg/ml adjusted to pH 7.4 and stored at 4°C until use. A second stock solution was prepared (0.25 mg/ml) then diluted to three final concentrations (25 µg 0.5 mg/kg, 50 µg, 1 mg/kg, 100 µg, 2 mg/kg). The lowest dose was administered by subcutaneous (s.c.) injections until P10, followed by intraperitoneal (ip) injections of the two higher doses with maturation. A schematic diagram of our subconvulsive treatment regimen is illustrated in **Figure 1**. Open handling was tested daily before the next dosing. Additional behavioral tests were performed on P21 and P22 and KA was re-injected at the end of behavioral testing to keep a 24 h interval between tests. All animals remained with their lactating mother and then sacrificed on P23, at 24 h after the last subconvulsive injection. There were 11 litters in this study, with approximately half treated with low doses of KA (n=58) and the other with PBS (n=54).

**Figure 1.**
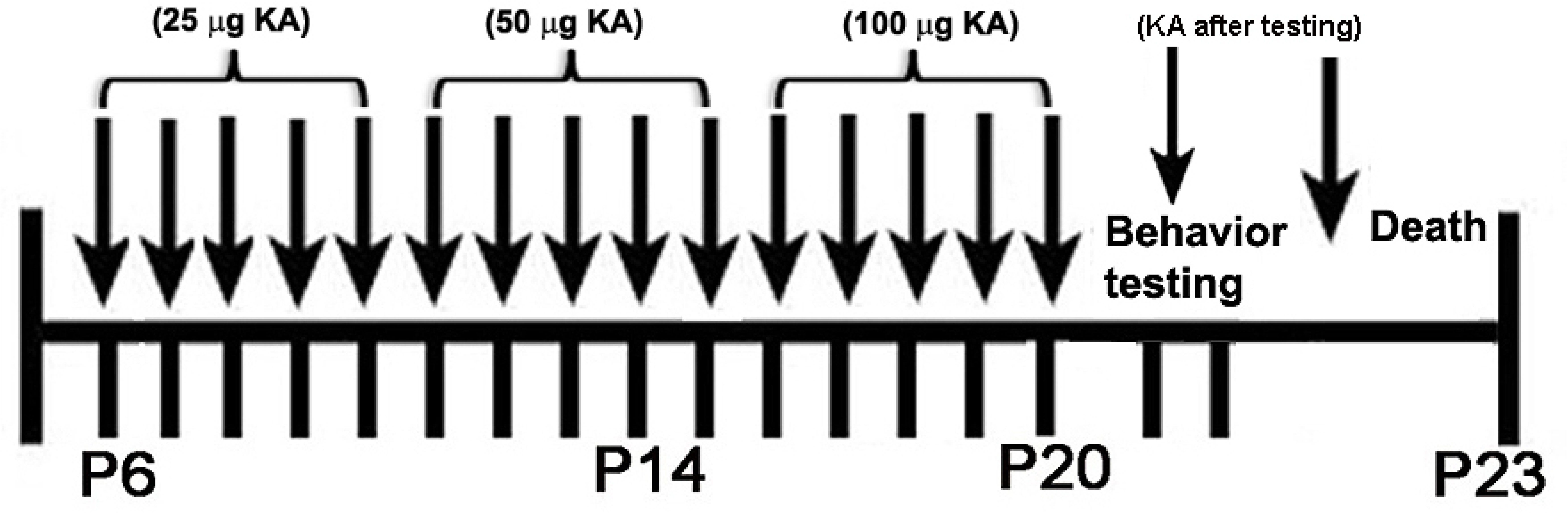
Schematic diagram of subconvulsive treatment regimen. Three subconvulsive step-up doses of KA were administered (25 μg, 50 μg, 100 μg) to rat pups for 15 days beginning on P6 with 5-day intervals. All Behavioral tests were carried out on P21. The animals were sacrificed on P22 or P23.

### 2.2 Open Palm Handling Test

A free to move around handling test was originated for the pre-juvenile ages of our study to monitor daily activity 24 hr after each subconvulsive treatment. This test also allowed us to observe whether any seizure behavior was present. In contrast to gentle neonatal handling to improve coping ability and reduce stress with stroking [36], control and experimental pups were free to move about within two open palms joined together for 1 min by a single investigator, illustrated in *Figure 2*. Animals were tested once daily beginning 48 h after initiation of the chronic subconvulsive KA treatment. Behavioral measures scored included: nose poking, attempts to escape, and percentage of animals escaping within the one-minute trial. Nose pokes were counted as light to moderate touches into the palms or fingers. Attempts to escape or climbing were scored when forelegs were both observed outside of the palms while standing on hind legs. Escapes were recorded when animals climbed out of the palms with all four limbs. Latency to escape was calculated at the time they escaped (1 min) which terminated the trial. Following the open handling testing, experimental pups were re-injected with micro-molar step up doses of KA while control pups were injected with PBS (see *Figure 1 schematic*).

**Figure 2.**
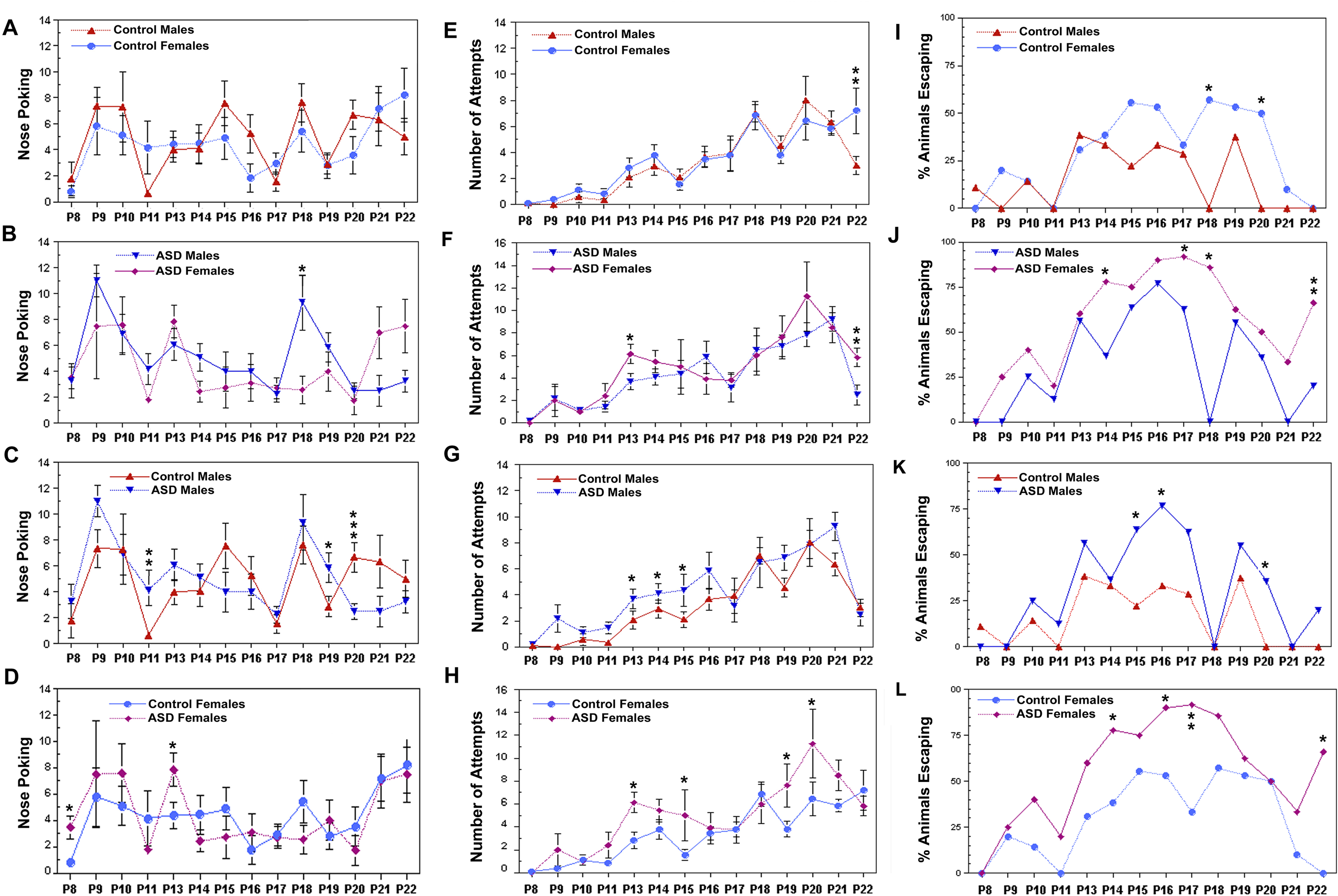
Sex-differences in the open handling test were observed. (**D**), Nose poking and (**B**-**H**) attempts to escape and (**I-L**), actual escape number and latency to escape were measured. (**A**), Nose poking in male vs. female controls was similar. (**B**), ASD males nose poke more often than ASD females. (**C**), There was an increase followed by a decrease in nose poking with age in the ASD males compared to male controls with maturation. (**D**), In contrast, ASD females had a higher rate of nose poking but only on P13 when compared to control females. (**E),** Control males and females had a similar pattern of attempts to escape except on P22 they had more attempts. (**F**), ASD males and ASD females also had a similar pattern of attempts to escape except on P22 they had more attempts. (**G**), ASD males had more attempts on P13-P15 than male controls but not afterwards. (**H**), ASD females had more attempts on P13, P15, P19 and P20. (**I**), Control males and females had similar amounts of escaping until P14, and then females showed higher escaping that varied. (**J**), ASD females escaped more often than ASD males except between P11-P13. (**K**), ASD males escaped more often than male controls on P13, P15, P17, P19, P20, P22. (**L**), ASD females escaped more often than control females beginning on P10 and escaped more frequently with age, except on P20. Both sexes escaped more often after treatment. (**A**-**H**), Data points represent the mean averages ± SEM of control males (n=16), control females (n=16), ASD males (n=20), ASD females (n=8). Student t-test, (**I**-**L**), Two proportion population z test was used. *, p < 0.05; **. p< 0.01.

### 2.3 Open-Field Test

Locomotion of the pre-juvenile rats was examined in the open field test (OFT) 24 hr after the last subconvulsive treatment of KA. Testing the animals was performed by placing the pups, one at a time, into the center of an open field without visual cues for 5 minutes. Area dimensions were 42.5×42.50 cm. The inside area of the arena was = 26.56×26.56 cm (39% of total area). Movements were automatically recorded with Activity Monitor program software, splitting movements between the inside area and periphery (Med-Associates, St Albans, VT, USA) as described previously [37]. The software program was set to monitor average velocity, distance traveled, the number of zone entries, time ambulant, counts and time of ambulation and stereotypic behaviors (e.g. grooming), time resting, vertical count, and time vertical. The chamber was cleaned with ethanol (70%) between animal trials.

### 2.4 Elevated Plus Maze

The elevated plus maze (EPM) was used to examine anxiety-like behavior and overall activity in response to the chronic subconvulsive treatment on P21 or P22 as described [37]. The elevated plus maze consisted of four arms 10×50 cm with two open arms without walls and two enclosed by high walls to block out the light. The entire apparatus was positioned 60 cm above the ground. The ends of the open arms were illuminated by lamps in order to further distinguish between them. Animals were tested one at a time for a 5 minute trial. The point of placement for each animal was in the center platform facing the open arm. Between each testing trial, the maze was cleaned with ethanol (70%), wiped with water and then toweled dry. Any-Maze software 4.3 was used to collect behavioral data for analyses. The following parameters were evaluated: number of entries into the closed vs. open arms (proximal and distal zones), distance traveled (m) within the closed and opened arms, average speed of subjects, time spent in the closed and opened arms, and total time active in the elevated plus maze.

### 2.5 Social Interaction: 3 Chamber Sociability and Novelty Test

The rectangular three-chambered described for mice was used to identify deficits in sociability in pre-juvenile rat pups as described [38]. Each chamber measured 20 cm (length) × 40.5 cm (width) × 22 cm (height). Dividing walls were made from clear Plexiglas, with small openings (10-cm width × 5-cm height) that allow access into each chamber. Testing of was conducted in three 5 min sessions, habituation, trial 1 and trial 2. The start location for each animal was in the center chamber that was left empty. A metal weight was placed on top of the wired chambers to prevent climbing. After habituation to the three empty chambers, an empty wired cup (novel object) was placed in the opposite chamber from the wired cup that trapped a never-before-met intruder pup of the same sex in trial 1. In the second trial, a different second never-before-met intruder pup was placed into the wired trapped cup whose location was swapped with the empty novelty wire cup from the previous trial, the “social novelty” session. Between each trial the three-chamber box was cleaned with ethanol. Any-Maze tracking software was used to analyze the time spent in each zone, the distance travelled, the number of entries to each zone, the number of visits to each zone, the number of line (zone) crossings, the time and number of mobile and immobile episodes, the time and number of freezing episodes, the latency to entry into each zone. Two-tailed Student’s t-test was used to statistically evaluate differences between control and ASD groups.

### 2.6 Group Social Interaction: Huddling Test Environments

Two litters of rat pups were simultaneously raised with their corresponding lactating mothers in separate cages in order to examine stranger to stranger interactions in a group setting. Huddling was used as an index of sociability. Control and treated pups were raised in the same cage. Five pups were treated with low subconvulsive doses of KA and five were treated with the same volume of PBS in two separate cages then tested separately. Three shapes of open environments with different sizes were used to examine group social behavior with siblings and non-siblings treated with PBS or the subconvulsive dose regimen of KA. The first environment was a small rectangular environment (34.29 x 43.18 cm). Five control mixed sex siblings were placed into the center together for a single trial without prior accommodation for 5 min. After cleaning the box with alcohol, five ASD siblings were placed into the arena for 5 min. Next, sibling and non-sibling pups of mixed sexes were re-tested in the same manner in a small circular environment similar to the size of rectangle environment (diameter = 43.18 cm). Last, sibling and non-sibling pups of mixed or unmixed sexes were introduced to a larger square environment (78.74 x 78.74 cm). Animals were videoed for 5 min trials. Grouping was counted from the videos which were analyzed manually every 5 seconds by an investigator blind to the experimental condition. Overall activity in the huddling tests was analyzed with Noldus Ethovison 13 XT software.

### 2.7 Object Recognition Test

The novel object recognition (NOR) test was used to analyze short-term and working memory of pre-juvenile rats subjected to chronic postnatal excitation compared to control baseline ability as described [39]. Rat pups were allowed to explore two identical objects for 5 minutes on day 1. Following a 24 h delay, on day 2, the animals were presented with two objects: one object was the same as in the first exploration trial and the second object was a novel object. The positions of the conditioned and novel objects were consistent between animals. Rodents have a tendency to interact more with a novel object than with a familiar object. The ability to store and remember information, or form memories, was measured by the rat’s ability to recall the conditioned object and spend more time with the novel object. If the animal did not spend more time with the novel object, then deficits in the ability to form memories was assumed.

### 2.8 Histology

Nissl staining with cresyl violet was carried out on fixed serial air-dried sections (40 μm) from serial vibratome sections from control and ASD groups. Histology included different levels of the hippocampus and amygdala along the septotemporal axis. Nissl labeled sections were mounted, processed through graded ethanol, and cleared with three changes of xylene for 15 min each. Sections were viewed and photographed under bright-field optics and phase contrast according to the rat atlas [40].

### 2.9 Statistics

Significant differences were determined for all groups for all behavioral tests. Results are given as means ± SEM. Values were subjected to Student’s test, One-way and Two-way analysis of variance (ANOVAs) using Excel and Sigma Stat software (Sigma Plot, 12.0, Software, Inc., CA). The Holm-Sidak post-test comparison factor was used for all pairwise multiple comparisons because this method is more powerful than the Tukey and Bonferonni tests for measuring differences between groups with unequal sample sizes. Escape was calculated as percent of animals escaping in the handling test and data points were analyzed with using the difference in population proportions. P values were calculated using z scores. Latency to huddling was analyzed with the survival analysis test with one censored variable, Mantel-Cox test for grouping time. Overall activity in the group huddling tests was also analyzed with Noldus Ethovison 13 XT software.

## 3. Results

### 3.1 Growth

Pups were weighed daily from P6 until P22. Chronic subconvulsive treatment with daily step up subconvulsive doses of KA had no effect on developmental weight gain between control and experimental groups. At 48 h after the beginning of treatment control males on P8 weighed nearly the same as the treated group (C: 18.5 ± 0.67 vs. ASD: 19.6 ± 0.8, N=17). Female weights were also not different on P8 (C: 14.7 ± 0.66 vs. ASD: 16.6 ± 0.74, N=9). Near the end of treatment before behavior testing on P19 there continued to be no difference in weight between control and experimental groups; control and ASD male weights grew at a similar rate (C: 44.9 ± 0.68 gr vs. ASD: 45.88 ± 1.58 gr, N=17). Although the females grew at a slower rate compared to the males, the treatment did not affect their normal growth patterns (C: 32.7 ± 2.0vs. ASD: 37.76 ± 3.2, N=9). The pups appeared to have normal dam-pup interaction while feeding in their home cage with the lactating mother.

### 3.2 Open Palm Handling Test

In contrast to calming handling tests used to reduce stress by gently holding the animals and stroking them for minutes or seconds daily [36, 41], a novel modified open palm handling test (with no restraining or stroking of the animal) was designed to test changes in daily behavior during the course of treatment as well as to see if any seizure behavior was present. The handling test served as a reliable index of progressive and cyclical behavior that varied according to sex and age. Beginning on P8, control and experimental animals were monitored 24 h after each KA injection, for 1 min trials. Three parameters were examined: nose poking, attempts to escape and percentage of animals escaping during the course of treatment. Daily poking activity was not different between control males and females throughout the study (**Figure 2A**). However, there was a decline in nose poking with maturation in both sexes of ASD pups due to spending more time climbing and escaping, but on P18 a significant increase in poking was observed in ASD males compared to ASD females (**Figure 2B**). In the ASD males, poking first increased then decreased with age more than male controls (**Figure 2C**). In contrast, ASD females only had a high rate of nose poking on P13 when compared to control females (**Figure 2D**). Both control males and control females exhibited a similar pattern of climbing attempts to escape with females escaping more often at P22 (**Figure 2E**). Sex differences in the number of attempts were observed only on P13 and P22 when comparing ASD males vs. ASD females (**Figure 2F**). In contrast, there was an increase in escape attempts on P13, P14, and P15 in ASD males compared to male controls (**Figure 2G**). For ASD females, attempts to escape were more cyclic being elevated on P13, P15, P19, and P20 (**Figure 2H**).

Sex differences also were seen with escaping. Control females escaped more often than control males between ages P15-P20 on an average of 33 ± 10% (**Figure 2I**). Similarly, ASD females escaped more often by 33.5 ± 9% compared to ASD males beginning on P14 which continued until the end of the chronic subconvulsive treatment (**Figure 2J**). When comparing only males, there was a transient period of a high percentage of escaping in the ASD animals from P15-P17 which was 40 ± 3.1% greater than the corresponding age-matched sibling controls. When comparing only females, a higher frequency of escaping was observed in ASD pups as early as P10, continued to P19, dropped to control values on P20, then increased again o P21 and P22 so that the average increase with maturation rose by 38 ± 1 7.3 % (**Figure 2L**). Therefore, the percentage of animals escaping increased in the 2^nd^ week of treatment when glutamatergic connections are maturing. There was no overt seizure behavior observed on any day during the handling test.

### 3.3 Open Field Test

Control and experimental groups were tested in an open field apparatus to quantify general differences in locomotor activity. At P21, weanling rats were placed into the center of the apparatus, one at a time, for 5 minutes without visual cues at 24 hours following the last subconvulsive KA treatment. Both ASD males and ASD females showed a significant increase in their average velocity when compared to PBS injected controls but not between each other (Figure 3A-C). Parameters such as total distance traveled, time spent ambulatory, and stereotypic behaviors were unchanged.

**Figure 3.**
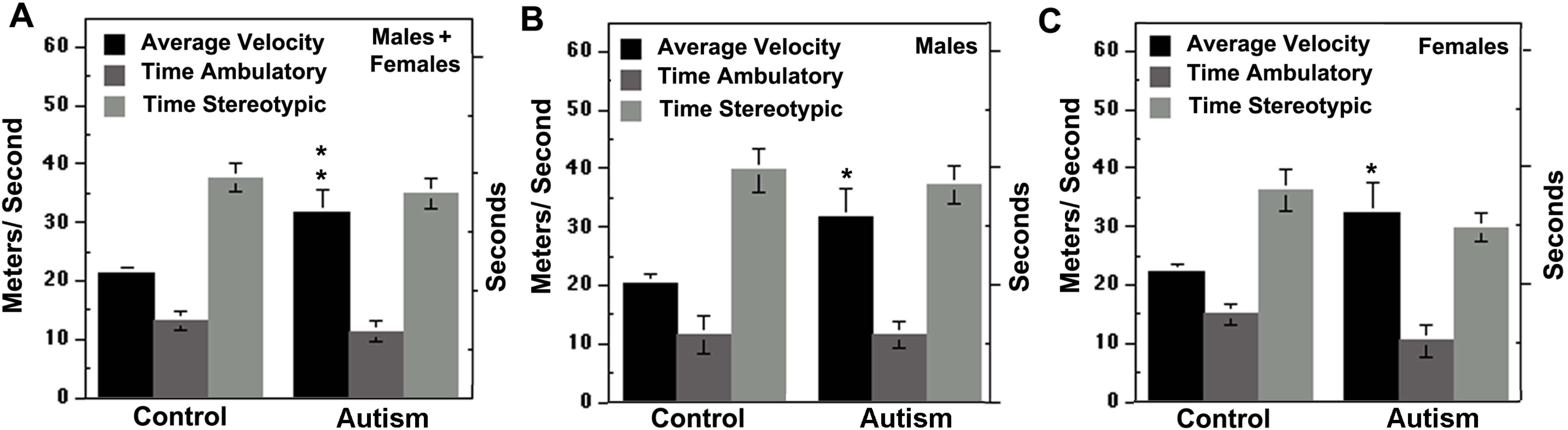
Open Field Test measured activity levels in male and female rats after exposure to subconvulsive KA. (**A**), Average velocity for the ASD group was faster than the control group with no significant changes observed in time ambulatory and time stereotypic. (**B**), Average velocity for was increased in ASD males and (**C**), in ASD females was compared to their respective sex matched controls. Bars represent the mean averages ± SEM of control males (n=13), control females (n=15), ASD males (n=17), ASD females (n=10). *, p < 0.05; **. p< 0.01, p< 0.001, Two-way ANOVA and Holm-Sidak multiple comparisons.

### 3.4 Elevated Plus Maze (EPM)

The EPM was used to examine sex-difference in anxiety-like behavior and overall activity in response to the chronic subconvulsive treatment. The total distance traveled in the EPM was greater in male controls compared to age-matched control females (p<0.001) (**Figure 4A**). After treatment, ASD males became more docile so that the total distance traveled compared to male controls was significantly reduced (p<0.05) whereas distance travelled by the ASD females was unaffected (**Figure 4A**). Similarly, the total time of activity in the EPM was greater in control males compared to control females. Following treatment, the opposite was observed; ASD males became less active and females more active revealing a significant interaction between sex and treatment (df = 3, F = 2.97, p<0.05) (**Figure 4B**). Accordingly, a significant interaction between sex and treatment was observed whereby the number of entries into the closed zones was greater for control males than for control females or ASD males but not ASD females (df = 3, F = 6.15, p = 0.002). Likewise, the number of entries into the closed arms was increased in ASD females compared to female controls and ASD males suggesting the females reach a higher level of anxiety at this developmental stage in this model (**Figure 4B**).

**Figure 4.**
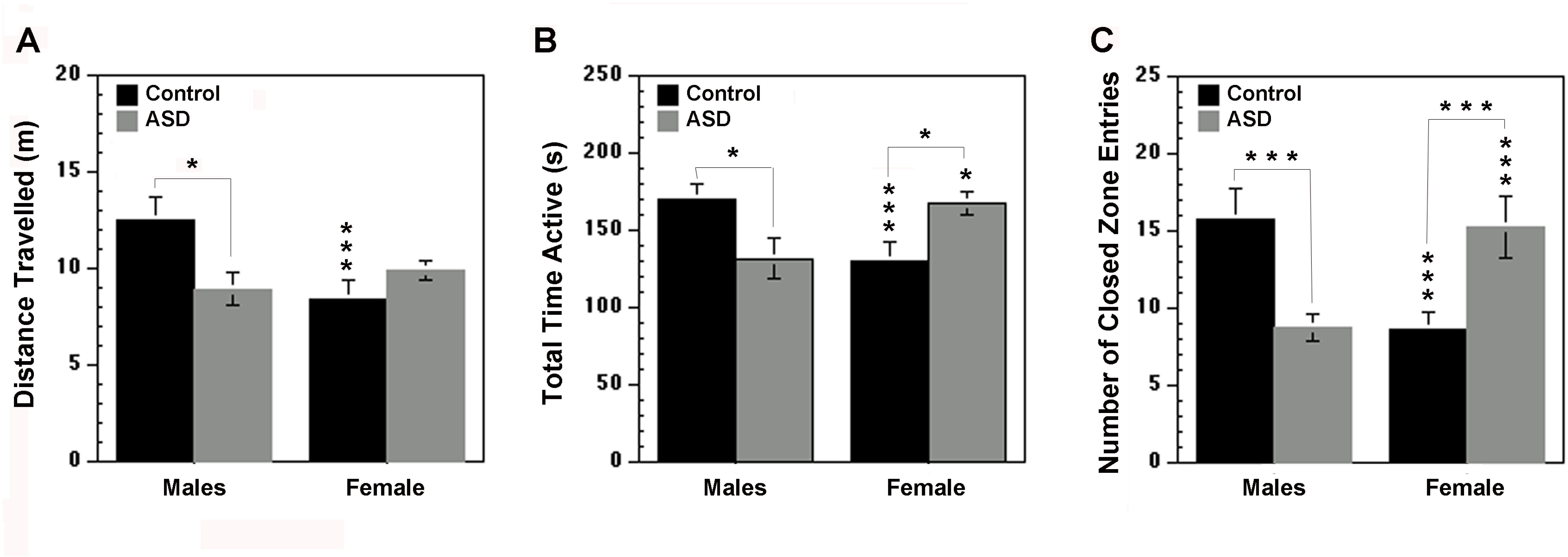
Elevated Plus Maze. (**A**), The total distance traveled was greater in control males compared with control females and ASD males, whereas ASD female distance traveled was unchanged. (**B**), The total time active decreased for ASD males but increased for ASD females compared to their respective sex matched controls, with control males starting off more active than control females. (**C**), The number of entries into the closed arms was reduced in ASD males but increased in ASD females compared to their respective sex-matched controls, with control males entering the closed arms more often than control females. Bars represent the mean averages ± SEM of control males (n=10), control females (n=12), ASD males (n=10), ASD females (n=8). *, p<0.05; **, p<0.01, two-way ANOVA and Holm-Sidak multiple comparisons.

### 3.5 Social Interaction: 3-chamber sociability test

To determine whether chronic hyperactivity in early life causes changes in social behavior in the pre-juvenile period, the 3-chamber sociability test was used where experimental and control rat pups were matched with a conspecific rat pup placed in an animal trap within the chamber 24 h after the last injection. The animals were placed, one at a time, into the center zone and allowed to explore and habituate to the new environment for 5 min. After all animals were habituated, trial 1 began with an empty animal trap in a subdivision of zone 1 and an animal trap holding a stranger pup in a subdivision of zone 3. At the end of trial 1, the empty animal trap and caged animal trap were switched. In trial 2, the animals were retested to determine how often and how long they spent with the conspecific pup in the animal trap. In trial 1, when averaging data from both genders together, ASD animals visited the novel object more often than controls (p< 0.05) (**Figure 5A**). However, when the sexes were analyzed separately to their sex-matched controls, it was found that only the males exhibited this behavior (p < 0.5). Accordingly, when ASD males were compared with ASD females, the males visited the novel object more frequently confirming that sex differences were due to the treatment (p=0.009) (**Figure 5B**). However, when male and female data were averaged, it showed that the ASD group as a whole spent more time exploring the novel object (C: 6 ± 0.7 vs. ASD: 9.48 ± 1.4 s, p<0.05). Male and female controls were not different from each other in their sociability behavior visiting the animal trap in trial 1.

**Figure 5.**
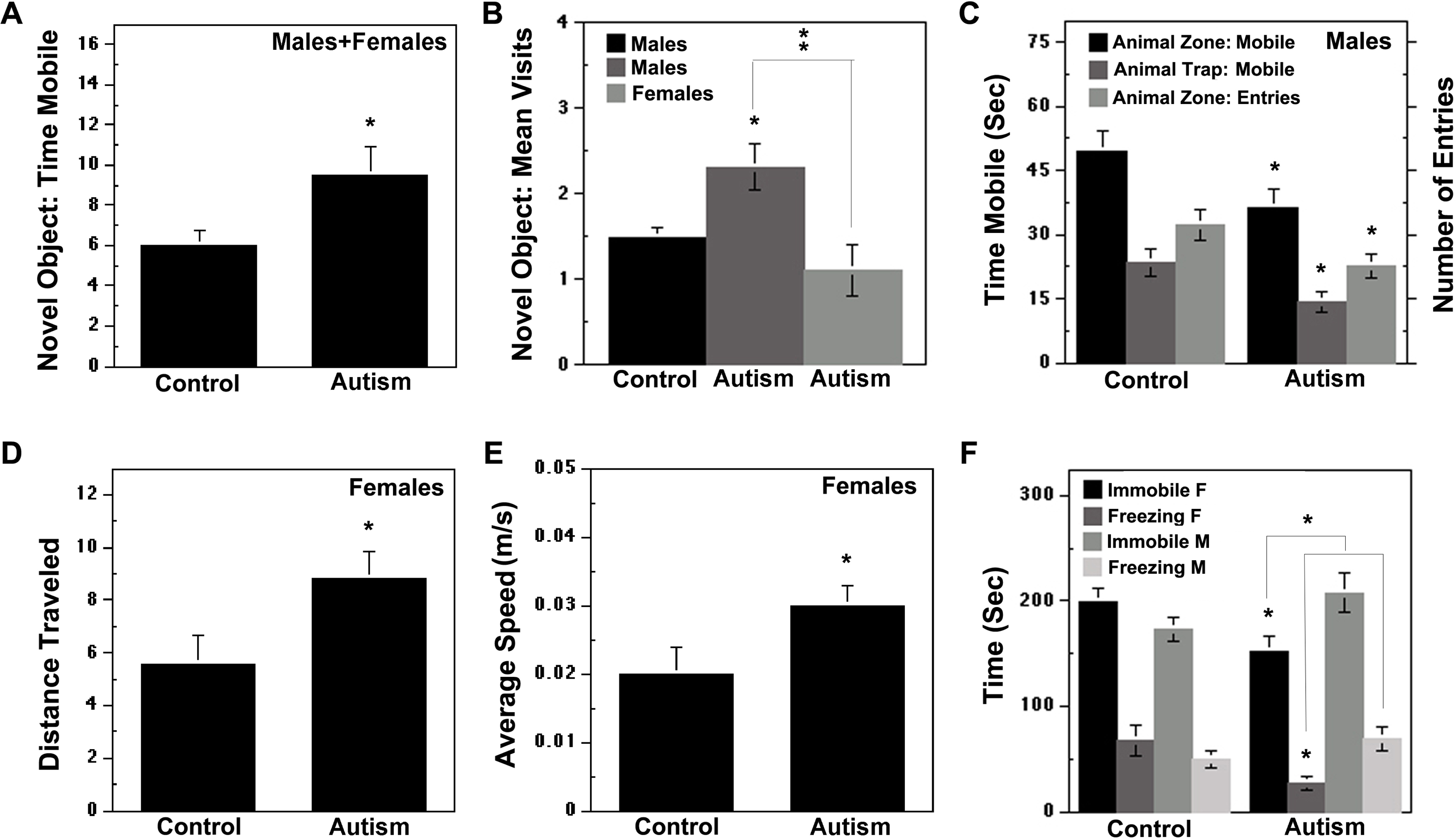
Social Interaction Test. (**A**), ASD rat pups spent more time exploring the novel object compared to control rat pups. (**B**), ASD males visited the novel object more often than both control males and ASD females. (**C**), ASD males spent less time exploring the animal zone and animal trap and visited the animal zone less often than control males. (**D**), Total distance travelled was greater in ASD females than control females. (**E**), ASD females had a faster average speed than control females. (**F**), ASD females spent less time immobile and freezing than control females, whereas ASD males spent more time immobile and freezing compared to ASD females. (**F**), No differences were observed in time spent immobile and freezing between control males and females. Bars represent the mean averages ± SEM of control males (n=12), control females n=17, ASD males n=16, ASD females n=10. Two-way ANOVA was used. *, p < 0.05; **. p< 0.01.

Main differences were observed in trial 2. Three parameters were reduced in the male ASD group: the number of entries in the zone containing the animal trap; the time mobile in the zone containing the animal trap; and the time spent with the animal (**Figure 5C**). In contrast, in the female ASD group, the mean speed and total distance traveled were significantly increased (Figures 5D-E). Moreover, ASD females were more mobile and displayed less time freezing compared to control females (**Figure 5G**). When ASD males were compared with ASD females, males were les mobile and spent more time freezing (**Figure 5G**). Male and female controls were not different in their sociability in trial 2.

### 3.6 Group Huddling Tests: 3 Environments

A novel group huddling test was used for social interactions in control and experimental animals. Sibling control and ASD pups were raised in the same cage until testing. Huddling was defined as 2 or more animals grouping very close together. Control and experimental groups of mixed sexes were tested separately from the same litter in each of the three environments for 5 min. Further testing was also done with non-sibling age-matched pups raised in a separate cage. The number of huddling events was determined by recording the number of animals grouping close together every 5 seconds over 5 minutes. Overall activity was also analyzed.

#### 3.6.1 Small Rectangular Environment

In the first experiment, animals were placed in a small rectangular environment (34.29 x 43.18 cm) for a single trial without prior accommodation. Control and ASD siblings were tested separately. Sibling trials consisted of either 5-7 pups per group that were allowed to freely move around and socialize within the small rectangle. Control siblings moved about the box in a non-anxious manner then quickly huddled together in a corner in groups of 5 or greater (**Figure 6A** and **C**). Control latency to the first grouping event minus one vs. total grouping occurred within 13.7 ± 4.3 sec vs. 77.5 ± 30 sec (p = 0.08). In contrast, ASD pups scattered about in the rectangular box in an anxious manner and grouped and huddled less often (**Figure 6B** and **C**). In ASD siblings, latency to grouping minus 1 and total grouping was significantly longer than controls (115 ± 55 sec vs. 280 ± 20 sec, F = 13.2 df, 13, p < 0.001).

**Figure 6.**
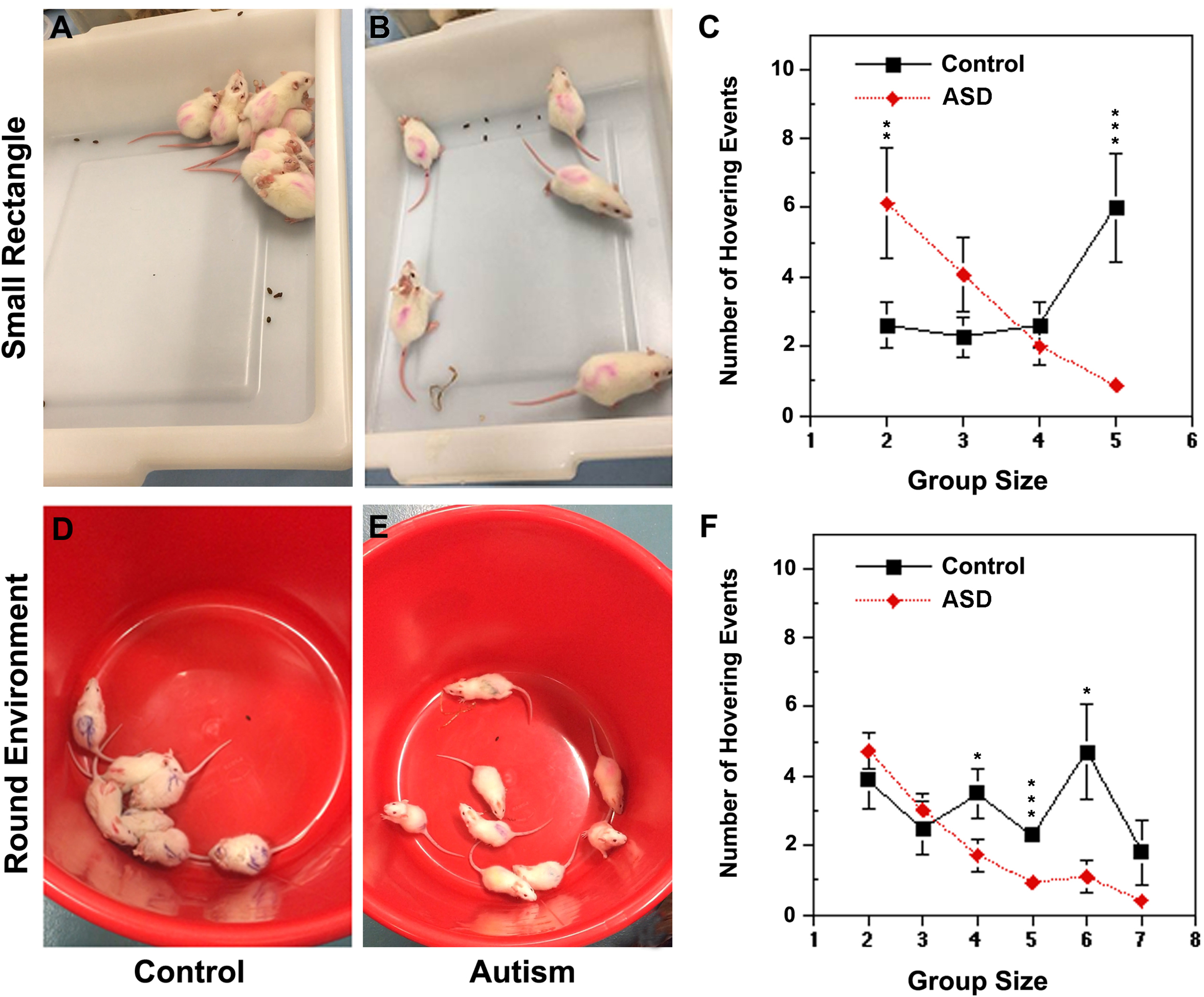
Huddling Test Rectangular and Circular Environments. (**A**), Control rat pups huddled in a large group in a corner of the small rectangular environment. (**B**), sibling ASD rat pups scattered about with less huddling. (**C**), Control animals huddled more frequently in larger groups of 5 compared to ASD animals, whereas the ASD animals huddled in smaller groups of 2 more often. (**D**), Control rat pups huddled in a large group in the round environment. (**E**), In the round environment, ASD rat pups scattered about and did not huddle in large groups. (**F**), In contrast, control animals huddled in groups of 4, 5 and 6 more often than ASD animals. (**C**, **F**), Data points represent the mean averages ± SEM of control of scatter plot graphs: (control, n=18; ASD:15, and control, n=13, ASD, n=19, respectively). Student’s t-test, *, p < 0.05; **. p< 0.01.

#### 3.6.2 Small Circular Environment

In the second experiment, pups were re-tested in a small circular environment similar to the rectangle environment (diameter = 43.18 cm). Sibling control animals quickly grouped to one side of the circle but were more spread out compared to the rectangular environment (**Figure 6D**). Quantifying the total number of huddling events showed that the controls grouped more frequently in greater numbers of 4, 5 and 6 compared to the ASD siblings throughout the testing period (t-test: p = 0.04, p = 0.005, p = 0.02, respectively) (Figure 6D-F). In contrast, the ASD sibling group scurried around the circular environment and huddled less frequently and in smaller groups (**Figure 6E** and **F**). When calculating total grouping −1 pup yielded the highest grouping with a 76.5% higher frequency rate in the control groups (p = 0.02). When non-sibling control groups entered the round environment, huddling still occurred more often and also in larger group sizes of 5 and 6 (p=0.04 and p=0.01) (not illustrated). In contrast, when two non-sibling ASD groups entering the round environment at the same time more circling and huddling in smaller groups compared to non-sibling controls was revealed (**Figure 6E** and **F**).

#### 3.6.3 Large Square Environment

In the third experiment, pups of mixed or unmixed sexes were introduced to the larger square environment (78.74 x 78.74 cm). When control siblings were tested from one litter, within the first 5 seconds they explored the box one by one or formed small groups followed by huddling into one corner (**Figure 7A**). In this experiment, the onset to initial huddling by the entire group occurred at 75 s (**Figure 7B**). The control pups remained in groups of 4 or 5 for the remainder of the trial. The litter matched ASD siblings (4 males/1 female) also roamed the large box but took longer time to huddle in a corner, at 105 s (Figure 7D-E). The control group spent 42% more time huddling all together compared to the ASD group that did not fully huddle. Accordingly, 3 averaged experiments showed the latency to the first total group huddling event was significantly shorter for the controls (45 ± 18 vs. 121 ± 9.6, p = 0.02). However, the number of huddling events per group size was not significantly different when sibling males and females were mixed (**Figure 7G**). When non-sibling control groups from different litters were introduced simultaneously in a new trial, huddling was quickly observed in large group sizes within 25 s in controls (**Figure 7C** and **H**). In contrast, when non-sibling ASD groups from different litters were introduced simultaneously, they ran around in a wild and erratic manner not observed in other trials. Huddling occurred in smaller group sizes and without corner specificity or preference (**Figure 7F** and **H**). Total group huddling was not observed in non-sibling ASD pups.

**Figure 7.**
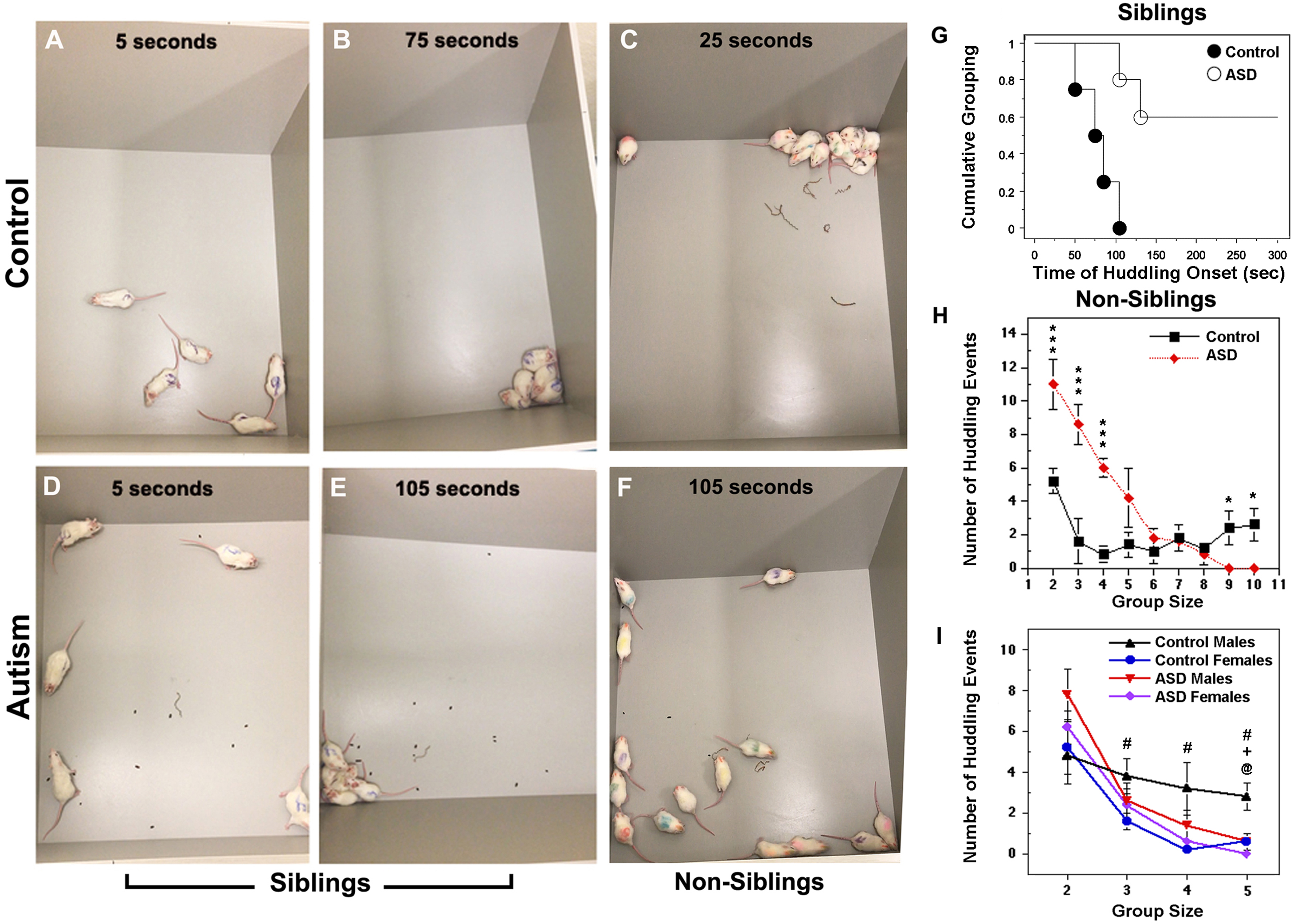
Huddling Test Large Square. (**A**, **B, D, E**), Control littermate siblings had a shorter latency to total group huddling compared to ASD siblings littermates (75 vs. 105 sec). (**C**, **F**), Control non-siblings huddled in larger groups of 9 or 10 compared to ASD non-siblings; ASD animals were erratic and scattered about in smaller groups of 2, 3 and 4 more often than controls. (**G**, **H, I**), Graphic analysis of huddling events of same and mixed litters. (**I**), When littermate sibling animals were tested by sex, control males huddled in larger groups and more frequently than control females. ASD males huddled less often in large group sizes compared to control males. ASD females huddled in smaller groups sizes similar to control females and ASD males, except that the entire group never fully huddled as did the ASD sibling males. (**G**), Kaplan-Meier Cum. Survival Plot software was used for grouping time, groups (control groups, N=4; ASD groups, N=4. (**H**), Data points represent the mean averages ± SEM of control, n=13; and ASD, n=14 animals. (**I**), Mean averages ± SEM of controls, n=10 and ASD, n=10). @ indicates significance between control males and ASD males, + indicates significance between control females and ASD females, # indicates significance between control males and control females, p<0.5, Student’s t-test.

The recordings of group huddling activity were further analyzed for mean overall activity as percent activity within the arena. The percent average activity in the arena of non-sibling ASD groups with males and females raised together in separate litters was increased by 169% compared to the control group (C: 0.842 vs. ASD: 0.312 mean % change in the arena). In a second experiment when males and females were raised in separate litters and then mixed in the arena, percent average activity within the arena was also elevated but to a lesser extent (C: 0.368 vs. ASD: 0.535 %). When the animals were tested by sex in this environment, the control males huddled in larger groups more frequently than the control females (**Figure 7I**). After treatment, ASD males huddled less often in large group sizes than control males. In contrast, ASD females huddled in smaller groups sizes similar to control females and ASD males except that the entire group never fully huddled as did the males.

### 3.7 Novel Object Recognition Test

The NOR test was utilized to evaluate the short-term and working memory of the ASD group as compared to control baseline levels. On day 1 during the conditioning trial, ASD males and ASD females explored the novel object to a similar degree as the matched control groups. On day 2 Sex differences were observed. Overall the control and experimental groups spent more time with the novel object compared to the conditioned object (Figure 8A-B). However, the ASD male rat pups spent much less time exploring both conditioned and unconditioned (novel) objects compared to control males, respectively (C:10.23 ± 2.84 vs. ASD:1.19 ± 0.56 s and C:14.13 ± 2.9 vs. 4.26 ± ASD:1.49 s, p<0.01) (Figures 8A-B). In contrast, ASD females did not significantly differ from the control females. There was a significant difference in sex and treatment but not an interaction between the two. Control females spent much more time exploring the novel object than control males (F: 34.2 ± 7.96 vs. M:14.13 ± 2.9 s, p<0.01) (Figure 8A-B). Females spent more time exploring the novel object (df = 1, F = 15.56, p<0.001) and the conditioned objects (df=1, F= 9.71 p=0.003) compared to control males and ASD males. While there was no sex difference in control mobility, there was an overall effect of treatment. The ASD males spent significantly less time mobile compared to controls (C:167.98 ± 14.93 vs. ASD:78.61 ± 13.51 s, (df = 1, F = 9.87, p = 0.003) (**Figure 8C**). Total time mobile for ASD females was greater than ASD males but did not differ from female controls (166.16 ± 21 vs. 78.61 ± 13.51 vs. 183.84 ± 18.72 s) (**Figure 8C**).

**Figure 8.**
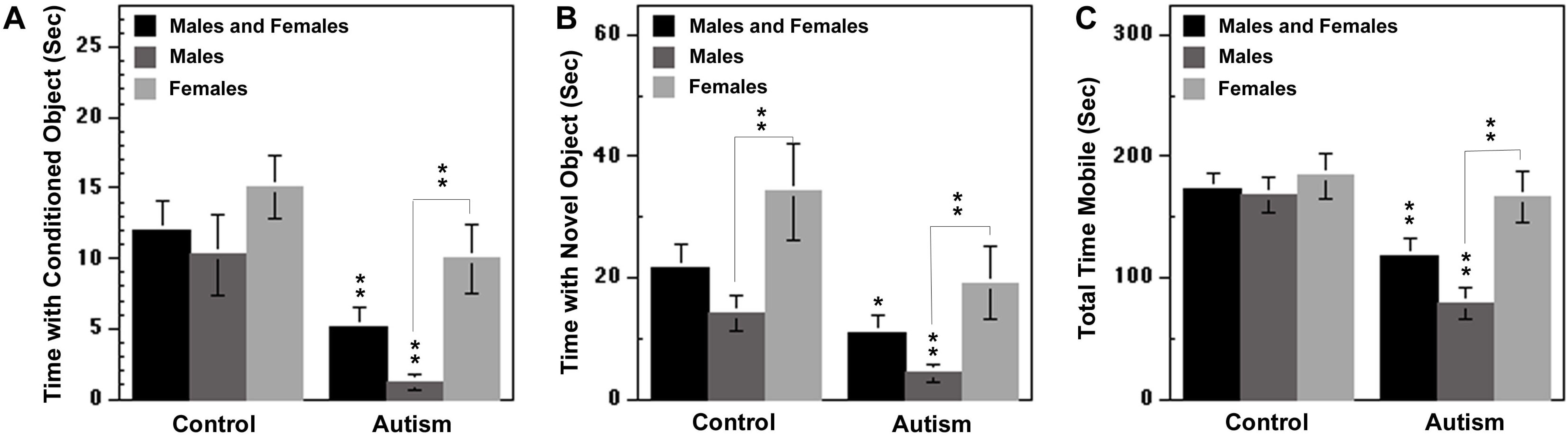
Novel Object Recognition Test. (**A**), Male and female controls spent similar time with the conditioned object whereas ASD male rat pups spent less time exploring the conditioned object. (**B**), Control females spent more time exploring the novel object than control males whereas, ASD male rat pups spent less time exploring the novel object compared to control males and ASD females. (**C**), Time mobile was similar for male and female controls. In contrast, ASD males were less mobile compared to controls and ASD females. Bars represent the mean averages ± SEM of control male (n=12), control female n=7, ASD male n=14, ASD female n=11. p<0.05; **. p<0.01, Two-way ANOVA and Holm-Sidak multiple comparisons.

### 3.8 Histology

Control and ASD coronal brain sections of the hippocampus and amygdala were stained with cresyl violet to estimate any changes in cellular morphology or neurotoxicity. In control hippocampus including the vulnerable CA1, staining was uniform and cell bodies were round and regular in shape (Figure 9A-B). In control amygdala and piriform cortex, staining was also uniform and cell bodies were round and regular in shape (Figure 9C-D). Similar to controls, staining of ASD brain sections was uniform and cell bodies were round and regular in shape throughout the hippocampus and amygdala with no obvious changes in cellular morphology or neurotoxicity (Figure 9E-H).

**Figure 9.**
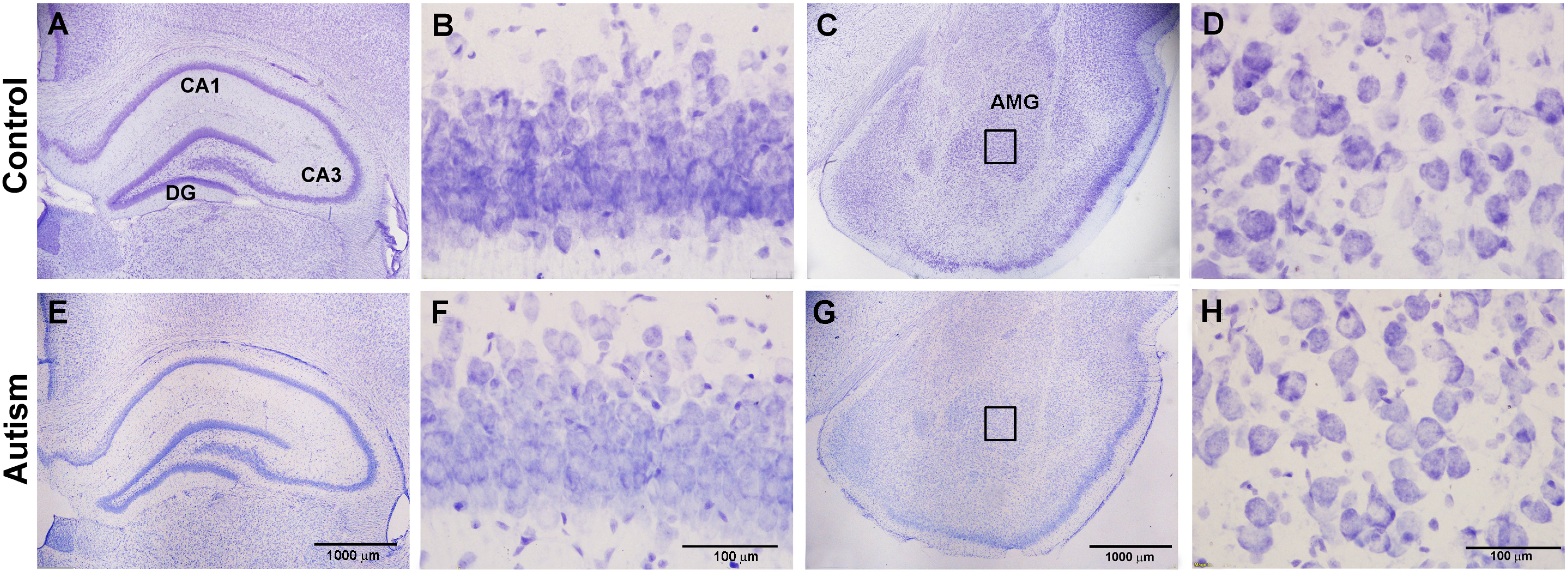
Photomicrographs of control and ASD coronal brain sections of the hippocampus and amygdala were stained with cresyl violet to estimate any changes in cellular morphology or neurotoxicity. (**A**), In control hippocampus, staining was uniform and areas were well defined (20x magnification). (**B**), At higher magnification (400x), cell bodies were round and regular in shape, illustrated for the CA1 subregion. (**C**), In control amygdala and piriform cortex staining was also uniform and areas were well defined. (**D**), Higher magnification (400x, *box* from panel C) also revealed round, healthy cell bodies. (**E**-**H**), Similar to controls, staining of ASD brain sections was uniform and cell bodies were round and regular in shape throughout the hippocampus, including the vulnerable CA1, and amygdala with no obvious changes in cell density, morphology, or neurotoxicity. Boxes in panels D and H reflect enlargements of basolateral amygdala in panels C and G, respectively. Scale bar = 100 μm or 1000 μm. **DG**, dentate gyrus, amygdala, **AMG**.

## 4. Discussion

The present study examined whether a subseizure postnatal state leads to mild forms of autistic pathology in a sex-related manner during the pre-juvenile and weanling period. Males were more susceptible to developing asocial behaviors and cognitive pathologies, whereas females were prone to higher levels of hyperactivity and anxiety. In our novel free to move handling test, ASD females escaped more often than ASD males. Accordingly, ASD females increased their mobility and number of entries into the closed arms of the EPM, consistent with both prenatal and postnatal VPA models of ASD [42, 43]. It appears that females may already be predisposed to a higher state of anxiety that was simply facilitated by the chronic subconvulsive treatment but insufficient to alter most of their behavior and cognition, whereas males with a lower baseline had less tolerability.

### 4.1 Social Behavior

In the 3-chamber social interaction test, sex differences were observed particularly in the second trial. The ASD males visited and spent more time with the novel object and the least amount of time and number of visits with the trapped stranger. Unexpectedly, ASD female social behavior under this setting did not differ from female controls despite their higher activity state. We noticed that the ASD males did not appear to sniff or interact with the stranger while spending time in the same chamber suggesting there was restricted interest not computed by our software program. This result, in part, contrasted findings reported in adult rats that had undergone one prior episode of KA-induced status epilepticus initiated on P7. As adults, males spent less time in the zone with the animal trap but more time sniffing the stranger rat compared to controls [44]. Differences may be due to developmental fluctuations in sex hormones such as follicular stimulating hormone (FSH), luteinizing hormone (LH), and testosterone levels in postnatal development [45].

Since the 3-chamber test is restricted to a 1:1 interaction, we sought further understanding of how overall activity and huddling behavior would be affected when the immature animals were placed with siblings or non-sibling peers in several novel environments. For example, in the largest novel environment (large square) the ASD siblings had a longer latency to initial group huddling compared to control siblings raised together. Non-sibling control pups also quickly huddled in large groups and remained in large groups throughout the trial, whereas ASD non-siblings did not huddle as one large group but rather moved around in an erratic fashion indicating less huddling and sociability occurred with strangers (non-sibling peers). Similar observations were observed in the round environment but to a lesser degree. Thus, ASD siblings were more social with each other than with non-sibling peers consistent with children that are better adjusted at home than in a public unfamiliar environment [46]. Huddling behavior during normal development in rodents depended on age-related activity of the somatosensory cortex and brain serotonin levels as well as increases in glutamatergic and GABAergic currents that abruptly occur during the second postnatal week, the time of our study [47]. This period also corresponds to the maturation of high affinity KA binding sites of the amygdala and its projections [48, 49]. However, the higher anxiety level in ASD females was not reflected as higher activity in the open field test or in group open huddling tests. This suggests that locomotion itself is not a core autistic trait in this model.

### 4.2 Glutamatergic Neurotransmission, KA and mGluR Receptors

Behavioral pathology may be due to altered activity within cortical and limbic areas of the amygdala complex due to the high density of high affinity of KA type receptors (KA1 and KA2) and low affinity glutamate receptors GluR5 and GluR6 which are expressed in these brain regions during early postnatal development [49–51]. In addition, dysfunctional group I metabotropic receptor (mGluR) mGluR5 has been observed in FRX mouse models which exhibit inappropriate trafficking that is associated with memory recognition deficits [52, 53]. Similarly, ASD male rat pups herein spent little time exploring conditioned and unconditioned objects; whereas ASD females did suggesting chronic subconvulsive activity treatment has sex-related effects on memory recognition that may involve post-synaptic mGluR5 receptors. Recently we observed that basal, unstimulated polyphosphoinositide (PI) hydrolysis was selectively reduced in the medial prefrontal cortex, thalamus, and amygdala following the chronic subconvulsive treatment (unpublished observation). Data suggest a deficit in slow G protein-coupled glutamatergic neurotransmission of the extra-hippocampal loop occurs in a subseizure model and could be responsible for deficits observed in the two-object recognition memory test and social interaction environments.

This subseizure model contrasts early life convulsive seizure models that over-excite glutamate ionotropic receptors and cause high elevations in glutamate and [Ca^2+^]_i_ which lead to concurrent epileptiform activity of the hippocampus and motor cortex as well as being associated with age- and region-dependent neurodegeneration [54–58]. Under stronger aberrant conditions a general decrease of KA and increase of NMDA receptor binding sites was previously observed in the hippocampus, somatosensory, piriform and entorhinal cortices regions involved with cognitive functioning [59]. Since NMDA receptors are critical for cognitive learning and neurological dysfunction, genetic studies identified missense mutations encoding the NMDA receptor GluN2B subunit that are associated with reductions in glutamate potency; increased receptor desensitization; and ablation of voltage-dependent Mg^2+^ block that may result in ASD phenotypes [60, 61]. Although not studied, these perturbations probably do not occur with the low doses of KA used. Histological evaluation revealed healthy appearing cells throughout the brain indicating that neuronal cell death was not responsible for the changes in behavior observed. Therefore, postnatal subseizure activity can lead to an ASD phenotype which is different and may be less severe from ASD phenotypes that result from early life convulsive seizures or genetic mutations.

Chronic postnatal increases in glutamatergic neurotransmission is also different from postnatal VPA-induced ASD since a single high dose of VPA, such as 400 mg/kg in neonatal mice, causes significant apoptosis in the cerebellum and dentate gyrus of the hippocampus [42]. Subconvulsive treatment also differs from a single VPA prenatal treatment administered on embryonic day 12.5 of pregnant rats since the offspring are not born normal and do not grow normally as they do here. Instead, they have lower body weight, delayed motor development, attenuated integration of a coordinated series of reflexes, and delayed nest-seeking responses [62]. Another difference is VPA is an anticonvulsant. It increases GABA concentration as well as having other mechanisms of action [63–65] making it difficult to decipher causes of milder forms of ASD. Micromolar doses of KA bind to high affinity KA binding sites to stimulate glutamate release. Retrospective receptor binding studies in adult rat forebrain tissues preparations revealed two apparent populations of KA binding sites exist in the nanomolar range: KD1 = 25-50 nM and KD2 = 3-14 nM [66, 67]. However, high affinity KA binding site distribution changes as a function of age with a transient but high expression in various brain stem nuclei and the stratum lacunosum moleculare of the CA1 area of the hippocampus at early postnatal ages [68]. In keeping, at P10 high affinity [3H] KA binding was observed in hippocampal preparations but not observed in the amygdala until P17-19, suggesting that certain aberrant behaviors would be expected to appear at different ages in response to chronic activation of KA-type glutamate receptors that may be reflected in the handling test. Accordingly, early postnatal administration of monosodium glutamate in rat pups during the first week of life caused elevations in the expression of low affinity KA receptor types, GluR5 and GluR6, high affinity KA1 and KA2 types, as well as [3H] KA binding at P21 which corresponds to the ages of our study [69]. In humans, distinct binding patterns of three main endogenous excitatory amino acid (EAA) receptor subtypes are well established early on by 20-21.5 gestational weeks suggesting over-excitation by environmental stress may begin in utero and continue ex utero, consistent with our postnatal rat pup model that reflects late *in utero* to neonatal periods to early childhood stages [70, 71].

### 4.3 Neurotransmission and Hypothalamic Pituitary Axis (HPA)

The excitatory/ inhibitory balance of ionic currents is also important for normal neural development [72]. In the mature CNS, activation of GABA_A_ receptor (GABA_A_R) Cl^−^ currents by GABA is the major means of inhibitory neurotransmission [73]. In development, inhibitory GABAergic signals shift from depolarizing to hyperpolarizing due to a reduction in intracellular chloride concentration ([Cl^−^]_i_) owing to the developmental regulation of NKCC1 and KCC2 sodium-potassium chloride transporter channel expression [74, 75]. This shift in GABAergic signaling is defective in genetic and prenatal animal models of ASD such as in FXS or animals treated with VPA *in utero* [76]. The chronic subconvulsive KA treatment herein was initiated postnatally when GABA has depolarizing properties and terminated during the weanling period (P20-P22) when GABA is inhibitory and there is a marked surge in glucocorticosteroids (CORT) [35, 71]. Chronic elevations of CORT can subsequently disrupt glutamate release and glutamatergic neurotransmission indirectly as well as GABAergic, cholinergic, noradrenergic and serotonergic and eCB systems leading to pathophysiology of mental disease [77, 78]. The age-dependent response to stress is also a critical factor for developing improved treatment strategies for ASD, as adolescent mice display a greater response to stress than adult mice due to prolonged activation of the HPA axis [78, 79]. Moreover, elevated levels of cortisol have been observed in ASD patients with callous-unemotional (CU) traits [80] as well as in patients with epilepsy which is further increased following seizures [81, 82]. Therefore, chronic elevations of basal glutamate in early life likely alter developmental sequences that induce aberrant circuit development and abnormal electrical properties of neurons that lead to autistic traits [76]. In keeping, acute hypoxia-induced neonatal seizures also result in long term increases in neuronal excitability, glutamatergic neurotransmission, and subsequent autistic-like behavior which was reversed by the mTORC1 inhibitor rapamycin when administered immediately before or after seizures [83]. This suggests that targeting the mTORC1 pathway signaling may also reverse milder forms of ASD elicited by subseizure activity. By the same premise, diminished stress-induced levels of cortisol have been observed in schizophrenia patients and other antisocial disorders such as early-onset conduct disorder [84, 85]. However, since not all autistic traits resemble schizophrenia, our data suggest that translational studies would be required to evaluate stress hormone levels in ASD patients as a function of age to determine applicable treatment.

### 4.4 Conclusion

Chronic subconvulsive activity in early postnatal development of naïve rat pups can lead to onset of ASD phenotypes in a sex-related manner in the juvenile weanling period. Males were more asocial and less cognitive or attentive while females were more hyperactive and anxious suggesting females may already be predisposed to a higher activity state which was facilitated by the chronic subconvulsive treatment. The type of test used may be important for early detection of ASD signs. Hence, treatment strategies to reverse adverse side effects of subconvulsive levels of glutamate may optimize an early clinical window of opportunity for clinical translation.

## Acknowledgements

We would like to thank our volunteer students Jessa Hofmann, Gabriella, Strzeletski, Pavankumar RamadassVenkatesulu, and Channa Wircberg who assisted with animal experiments and analytical aspects of the work. We also give many thanks to Rose Friedman and honor the memory of Marvin Friedman for providing seed money for the study.

